# A single giant cell at the origin of an evolutionary innovation

**DOI:** 10.1101/2024.02.08.579433

**Authors:** Arnaud Badiane, Séverine Viala, David Armisen, Ilona Mignerey, Christelle Forcet-Vauchel, Christina N. De Jesús Villanueva, Steven Van Belleghem, Antonin Crumière, Emilia Santos, Mirko Francesconi, Abderrahman Khila

## Abstract

Key innovations play a central role in the diversification of lineages, yet our understanding of the mechanisms underlying their emergence remains fragmented. The propelling fan is a key innovation associated with the diversification of *Rhagovelia* water striders into fast flowing streams. Here, we show that a single giant cell is responsible for the development of the fan throughout embryogenesis. RNA interference against the fan-specific gene family *gsha*/*mogsha* depleted the fan in the embryo but did not alter the giant cell, indicating that these genes are not required for specifying the identity of this cell. *gsha*/*mogsha* instead seem to activate the expression of cuticular genes during early development, suggesting a role of these genes in regulating the accumulation of fan structural proteins. We also show that Hox genes act to block the giant cell in the fore- and rear-legs, thus restricting its fate to the mid-legs – the pair of legs that generates movement on water. In addition, Hox expression is specifically excluded from the giant cell, thus allowing the expression of the fan’s developmental genetic program. This study reveals that a single cell can be sufficient to generate innovative traits that have been central for a lineage to acquire new ecological opportunities and burst into diversification.

**One-sentence summary:** A single giant cell orchestrates the developmental genetic program of a key innovation

## Main text

Key innovations are derived organismal features that permit a species to occupy previously inaccessible ecological states (Miller et al. 2022). From the pharyngeal jaws of cichlid fishes (Liem 1973) to the adhesive toepads of anole lizards (Miller and Stroud 2022), examples of how these new traits have bolstered diversity across the tree of life abound (e.g., (Siqueira et al. 2020; Garcia-Porta et al. 2022). Despite a long-standing interest in key innovations (Simpson 1953; Losos 2009; Erwin 2021), retracing their origin remains a major challenge for evolutionists, one that requires a multidisciplinary approach to investigate processes across biological scales (Wagner and Lynch 2010; Miller et al. 2022). Current evidence suggests that taxon-restricted genes play a major role in the origination of these new phenotypes (Khalturin et al. 2009), but how these genes are integrated within pre-existing developmental programs and how these programs alter the cellular composition underlying the trait remains unclear.

Water striders evolved a series of key innovations that allowed them to conquer the water-air interface (Andersen 1982; Hu et al. 2003; Crumière et al. 2016; Armisén et al. 2018). The genus *Rhagovelia*, also known as riffle bugs, experience an additional transition from stagnant or clam waters to occupy fast flowing streams – an ecological niche that is inaccessible to the extreme majority of water striders (Andersen 1982; Polhemus and Polhemus 2008; Armisén et al. 2022). This new ecological transition is associated with an impressive burst of diversification, such that the genus *Rhagovelia* alone accounts for over half of the family Veliidae and more than 20% of all water striders (Polhemus and Polhemus 2008). This diversification is associated with a unique key innovation consisting of a pair of plumy fans, of cuticular composition, located at the tip of the mid-legs, which are responsible for generating movement on the water. The fan, present in all *Rhagovelia* species, allows these animals to sustain locomotion against water current (Santos et al. 2017). Biomechanical examination of fan function revealed that the fan pierces the water while the tarsus remains at the surface to prevent the leg from wetting (Pennisi 2024). We previously found that two new paralogs, one named *geisha* (*gsha*) exclusively found in the genus *Rhagovelia* and the second, *mother-of-geisha* (*mogsha*) found in the Hemiptera, were required for fan formation during post-embryonic development (Santos et al. 2017). This discovery established that the emergence of new genes can contribute to the emergence of new traits that allow lineages to acquire hitherto inaccessible ecological opportunities. Here we sought to determine the cellular composition of the fan and the genetic interactions underlying its development in time and space during embryogenesis.

## Results

Previously, a visual color-based *in situ* hybridization screen identified five transcripts that are expressed in a cellular domain of the mid-leg which prefigures fan formation (Santos et al. 2017). To better characterize the cellular composition of this domain, we developed fluorescent *in situ* hybridization chain reaction (HCR-FISH) and used a *gsha/mogsha* probe as a fan-specific marker in the embryo. Surprisingly, this method revealed that *gsha/mogsha* expression was restricted to a single giant cell located at the tip of T2 embryonic legs, instead of a cluster of cells as previously thought. A time series using staged embryos revealed that the first signs of *gsha/mogsha* expression appeared at 96h after egg laying (AEL) in two (64%, n=9) to three (36%, n=5) cells located at the tip of T2 legs (Figure 1A). At 120h of development, all embryos had *gsha* expression in a single larger cell (∼30 μm of diameter) with a large nucleus suggesting that it might be polyploid (Figure 1B). By 144h, the cell became elongated along the proximal-distal axis and reached about 70 μm in length. At this stage, the distal pole of the cell is pointing out of the leg epithelium toward the position of the future fan (Figure 1C). At 312h, which marks the end of embryogenesis, the length of the cell is over 200 μm and its distal pole is connected through a cytoplasmic extension to two fork-like structures in the middle of which the fan has developed (Figure 1D). No *gsha/mogsha* expression was detected in T1 or T3 legs, consistent with the absence of the fan in these legs (Figure 1E) and confirming previous observations (Santos et al. 2017). These findings suggest that a single giant cell is at the origin of fan development.

**Figure 1.**
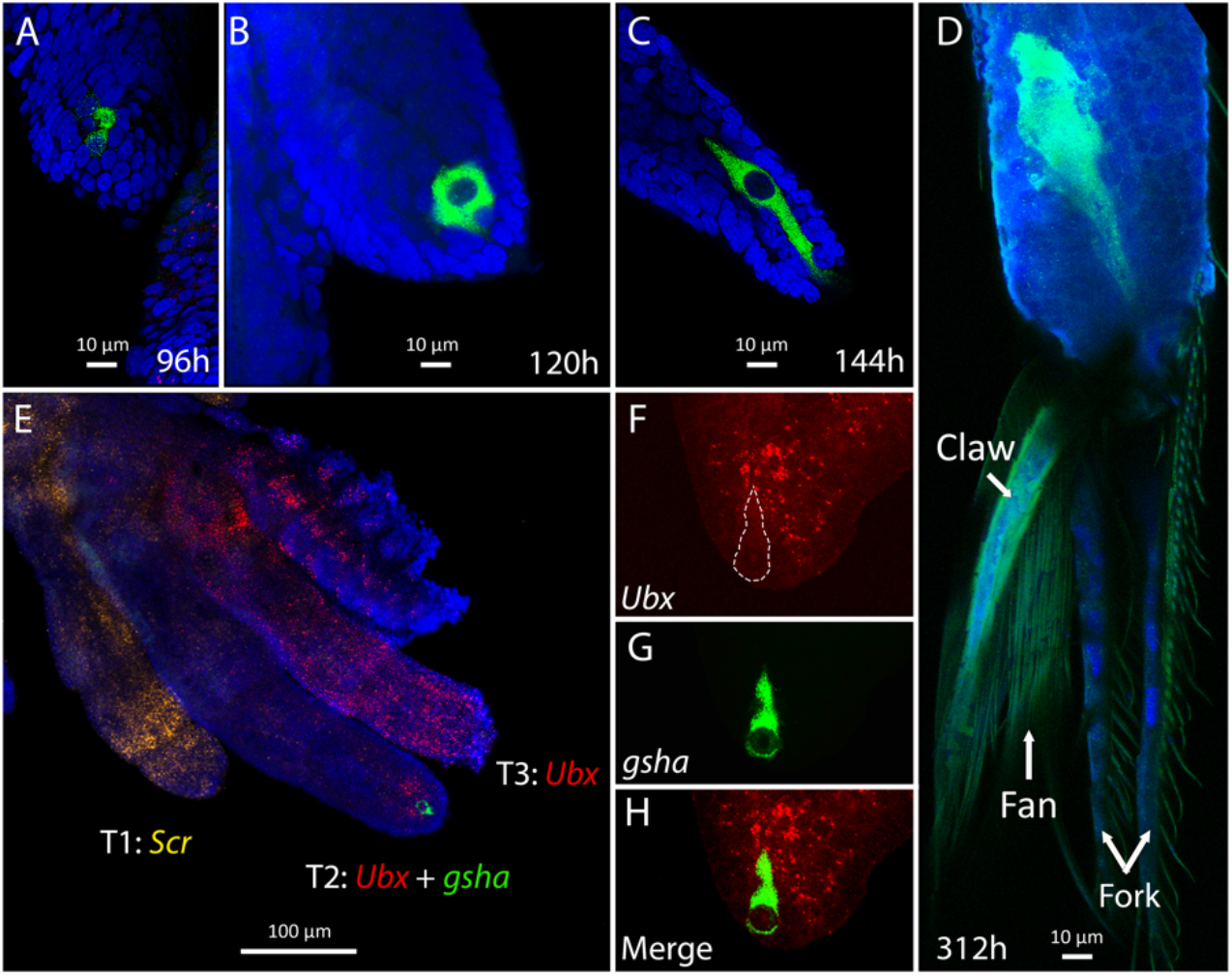
HCR-FISH images of *R. antilleana* embryos. Developmental time series of T2 legs at 96h (A), 120h (B), 144 (C) and 312h (D) after egg laying. We observe that the expression of *gsha*/mo*gsha* (referred as to *gsha* on the figure for the sake of brevity) is located in 2-3 cells at 96h, and is then restricted to a single cell that keeps growing throughout embryogenesis to become giant. At 312h (D), the embryo is fully developed and we can see the claw and the fan by autofluorescence, as well as the the two protrusions of the fork between which the fan folds and unfolds. Panel E represents the left side of an embryos with the T1 leg showing *Scr* expression, the T3 leg showing *Ubx* expression, and the T2 leg showing some Ubx expression and *gsha*/*mogsha* expression confined to the giant cell. We then show the tip of a T2 leg showing either only *Ubx* expression with the giant cell delimited by a dotted line (F), only *gsha*/*mogsha* expression (G), or both *Ubx* and *gsha*/*mogsha* expression (H) to demonstrate that *Ubx* expression is excluded from the giant cell.

Based on sequence similarity and the absence of introns in *gsha*, the gene *gsha* is thougth to originate from a retrotransposition of its ancestral paralog *mogsha* (Santos et al. 2017). However, the lack of a genome assembly prevented the annotation of their genomic architecture and a firm confirmation of this conclusion. A high-quality chromosome-level genome assembly of *R. antilleana* genome allowed the confirmation of the ancestral sequence *mogsha*, which is composed of four permanent exons and one alternative exon interspersed with introns (Figure 2). Surprisingly, the genome assembly revealed the presence of two copies of *gsha*, which we named *gsha1* and *gsha2*. Both *gsha* genes are intronless and are composed of a unique exon made of the juxtaposition of the four *mogsha* permanent exons (Figure 2). *mogsha* is located on a different scaffold than the two paralogs *gsha1* and *gsha2*, confirming our conclusion of the two new copies originating from a retrotransposition. Moreover, *gsha1* and *gsha2* are highly similar in sequence (97.29%), with *gsha1* being closer in sequence to *mogsha* (86.45% identity) than *gsha2* (85.37% identity). These two new genes are separated by only 2875 base pairs, suggesting that they could have originated from a tandem duplication of the new gene.

**Figure 2.**
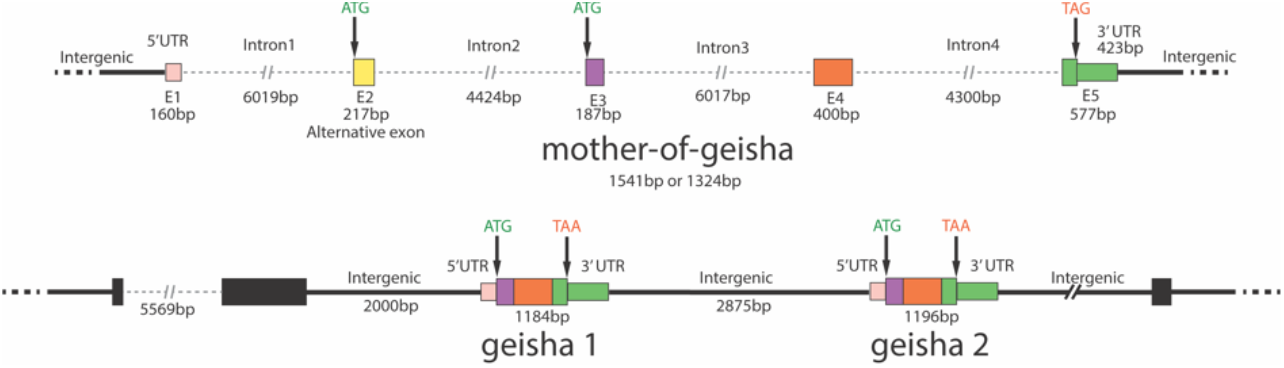
Representation of the genomic architecture of the gene *mogsha* (top panel) and its two paralogs *geisha1* and *geisha2* (bottom panel). The different colors in *gsha1* and *gsha2* represent the sequences that are homologous to the *mogsha* exons of the same colors. Exon 2 from *mogsha* is absent from both *gsha1* and *gsha2*. The black rectangles on the bottom panel represent sequences matching those of transposons.

Because the giant cell and the fan it produces are restricted to T2 legs we hypothesized that they may be regulated through the action of Hox proteins, which are known to establish the identity of the segments along the anterior-posterior axis. As in other water striders, the Hox gene *Scr* is expressed in T1 and *Ubx* in both T2 and T3 (Khila et al. 2009; Crumière et al. 2019). The domains of expression *Scr* and *Ubx* cover the distal tip of T1 and T3 legs respectively (Figure 1E). Interestingly, *Ubx* in T2 is excluded from the giant cell while being expressed in some of its surrounding cells (Figure 1F-H). These observations led to the hypothesis that Hox genes repress the formation of the giant cell and the fan. RNAi against *Scr* or *Ubx* induced the specification of an ectopic giant cell in T1 and T3 legs, respectively (Figure 3A, C). In these transformed legs, the giant cell expressed the fan-specific markers *mogsha*/*gsha1*/*gsha2* and *CP19*, and produced ectopic fans, confirming the role of these genes in blocking fan formation in T1 and T3 (Figure 3A, C). In T2 legs however, *Ubx* RNAi transformed the identity of the ventral claw but did not affect the giant cell or the fan (Figure 3B). This result is consistent with the absence of expression of *Ubx* from the fan cell and indicates that an unknown genetic factor blocks the expression of *Ubx* in the giant cell to allow fan development (Figure 1F-H). The induction of ectopic fans and transformation of the ventral claw were identical for *Ubx* and *Scr* knockdown for both parental and nymphal RNAi (Supp Figure S1), suggesting that the fan genetic program persists throughout post-embryonic development. These results suggest that the fan developmental program may have originated from the modification of the program that specifies the arolium and evolved as part of the ground plan of *Rhagovelia*. Hox genes act to block this program in T1 and T3 and the ability of T2 to activate the program required the exclusion of Hox genes from the cell that produces the fan.

**Figure 3.**
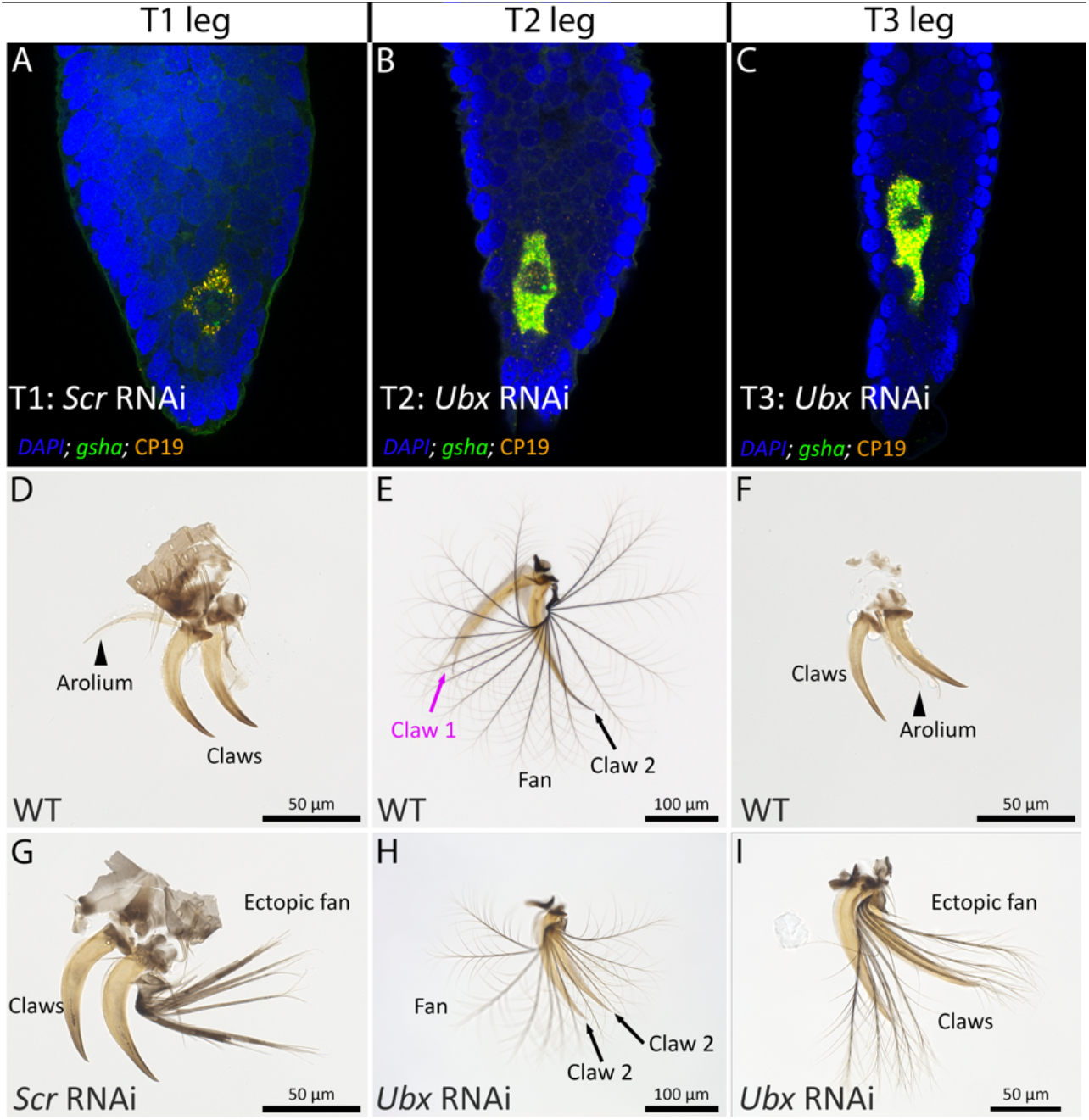
Images of embryonic and nymphal (1^st^ instar) phenotypes in *Scr*- and *Ubx*-RNAi individuals. In *Scr*-RNAi embryos, T1 legs acquire the giant cell that expresses *mogsha*/*gsha1*/*gsha2* and CP19 (A) which is normally restricted to T2 legs. When these embryos hatch, T1 legs which normally present an arolium (D) instead develop an ectopic fan (G). In *Ubx*-RNAi embryos, T2 legs still have the giant cell and its co-expressed markers *mogsha*/*gsha1*/*gsha2* and CP19 (B). Fan integrity is conserved in the hatching nymphs but the ventral claw loses its identity (E) in *Ubx*-RNAi embryos (H).

The genes *mogsha*/*gsha1*/*gsha2* are required for the development of the fan (Santos et al. 2017), but their position in the fan specification genetic program is unknown. These novel genes could either specify the identity of the fan cell by inducing cell fusion, endoreplication or growth, by playing a regulatory role that ensures the exclusion of *Ubx* expression from the fan cell, or by activating other genes required for fan formation. To test these hypotheses, we knocked down *mogsha*/*gsha1*/*gsha2* from the fan cell using RNAi and examined the expression of possible target genes using HCR. This parental RNAi experiment resulted in a severe depletion of the fan (Figure 4A-B) that is identical to the result we previously reported with nymphal RNAi (Santos et al. 2017). First, *mogsha*/*gsha1*/*gsha2* RNAi did not induce supernumerary fan cells or alter its shape or size (Figure 4I-N). Furthermore, *mogsha*/*gsha1*/*gsha2* RNAi did not induce *Ubx* expression in the fan cell (Figure 4I-N). These results exclude a role of these genes in specifying the identity of the fan cell and in repressing *Ubx* in this cell. Next, we tested the expression of two cuticular markers that are co-expressed with *mogsha*/*gsha1*/*gsha2* in the fan cell. When *mogsha*/*gsha1*/*gsha2* are depleted, *CP19* expression is strongly reduced in early embryos (Figure 4C-E, I-K). However, in late embryos, *CP19* expression is unaffected even when *mogsha*/*gsha1*/*gsha2* expression has been entirely erased (Figure 4F-H, L-N).

**Figure 4.**
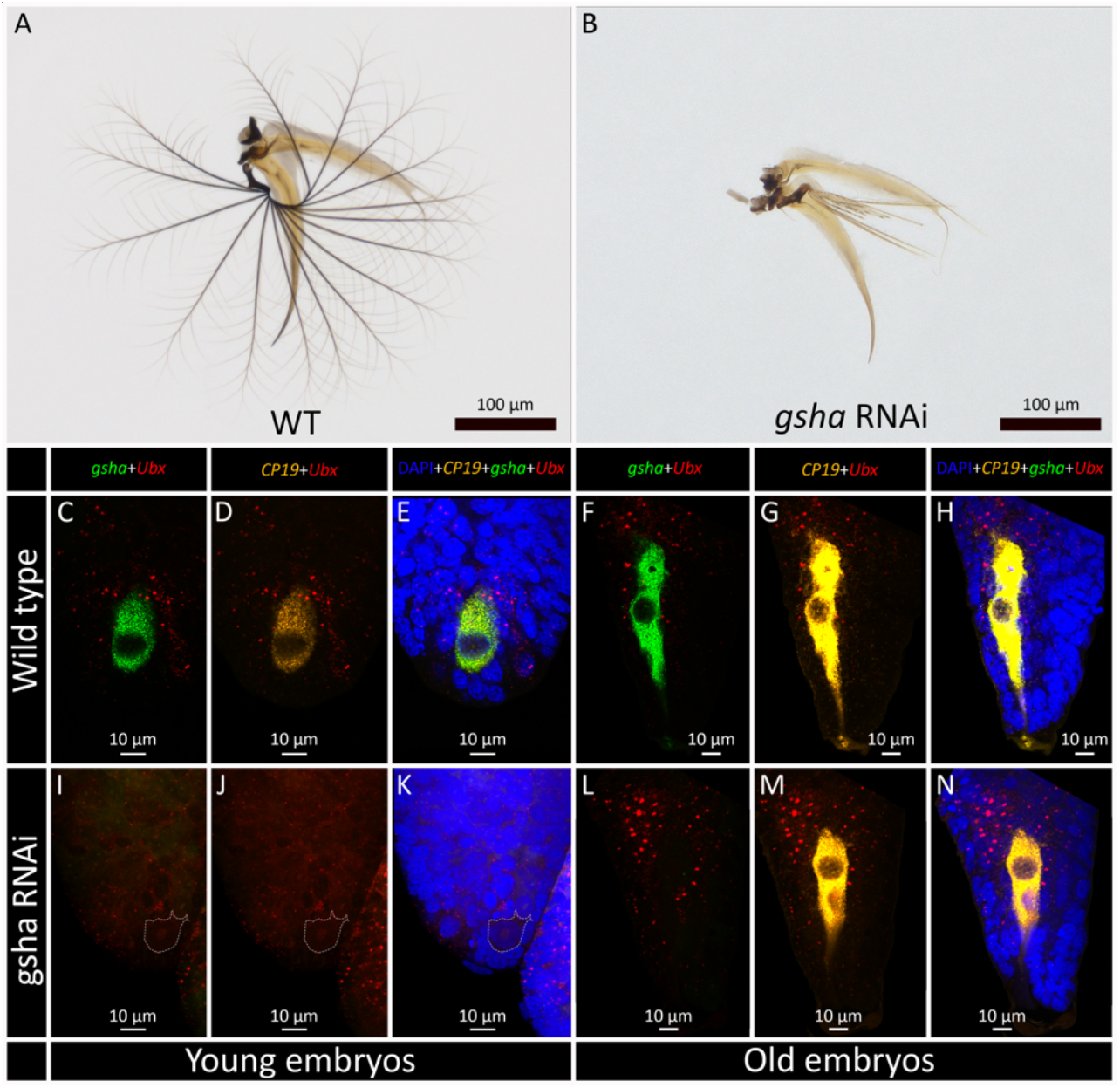
Images of *mogsha*/*gsha1*/*gsha2*-RNAi embryos. The knockdown of *mogsha*/*gsha1*/*gsha2* in embryos modified the wild-type fan (A) into a strongly impaired fan (B). HCR imaging of wild-type T2 legs in young embryos (before 144h) and old embryos (after 144h) are shown either with *mogsha*/*gsha1*/*gsha2* and *Ubx* expression (C, F), with *mogsha*/*gsha1*/*gsha2* and CP19 expression (D, G), or with the multiplexed expression of *mogsha*/*gsha1*/*gsha2*, CP19, and UBx, along with DAPI nuclei-staining (E, H). HCR imaging of *mogsha*/*gsha1*/*gsha2*-RNAi T2 legs in young embryos (before 144h) and old embryos (after 144h) are shown either with *mogsha*/*gsha1*/*gsha2* and *Ubx* expression (I, L), with *mogsha*/*gsha1*/*gsha2* and CP19 expression (J, M), or with the multiplexed expression of *mogsha*/*gsha1*/*gsha2*, CP19, and UBx, along with DAPI nuclei-staining (K, N). In young RNAi embryos, the cell is delimited by a dotted line. *CP19* expression is lost in young *mogsha*/*gsha1*/gsha2-RNAi embryos but is rescued in old embryos.

## Discussion

Here, we showed that the emergence of the propelling fan as a key innovation in the *Rhagovelia* lineage is accompanied by new processes at the molecular and cellular level. At the cellular level, a single giant cell is driving fan development. This cell also constitutes the exclusive expression domain of *mogsha*/*gsha1*/*gsha2* and *CP19*, and possibly other genes not targeted here. We indeed previously showed that gene expression in T2 legs was enriched with cuticular genes that likely contribute to fan formation (Santos et al. 2017). Furthermore, the cell acquires a peculiar teardrop shape throughout embryogenesis, with a cytoplasmic extension pointing towards the tip of the leg, precisely where the fan is attached to the leg. This supports the conclusion that the giant cell produces and transports most of the material (e.g., cuticular proteins) necessary to build the fan.

In early developmental stages, there are actually two to three cells that express *mogsha*/*gsha1*/*gsha2* which become a unique giant cell with a large nucleus later on, suggesting polyploidy (Figure 1A, B). Several mechanisms could explain this transition from several normal-sized cells to one that is very large. First, lateral inhibition, the capacity of a cell to inhibit the activity of its neighboring cells, has been described in neural and sensory cells, for example during the development of sensory bristle in *Drosophila* (Simpson 1990; Axelrod 2010). Under this scenario, only one of the 2-3 cells expressing *mogsha*/*gsha1*/*gsha2* at 96h would develop into the giant cell by inactivating the other two, and then undergo endoreplication to reach polyploidy (Lee et al. 2009). Alternatively, mechanisms of cell fusion are well known to give rise to large polyploid cells and to contribute to cell differentiation (Ogle et al. 2005; Losick et al. 2013; Orr-Weaver 2015). In both cases, given that the large nucleus and the high activity of the giant cell, we hypothesized that it might be polyploid. Polyploidization is known to be a way to increase cell size and gene expression (Frawley and Orr-Weaver 2015). Developmental growth by polyploidization rather than by increase in cell number can be advantageous because cell division cycles can impair some specific functions and may also disrupt barrier functions (Frawley and Orr-Weaver 2015). If the giant cell is not polyploid, it might be that the cell, which does not divide throughout embryogenesis, has its cycle fixed at the interphase, when the nucleus can double in size (Maeshima et al. 2011). In all cases, the giant size of the cell and its nucleus reflects a high transcriptional activity. Altogether, these results suggest that this cell may be a new cell type which is at the origin of fan development.

At the genomic level, we detected the presence of three paralogs (*mogsha*/*gsha1*/*gsha2*). This genomic architecture suggests that *gsha1* originated as a retrotransposition of *mogsha*, thus explaining the long distance between these genes which are located on two different scaffolds (i.e., two different chromosomes), and the intronless sequence of *gsha1*. Later, in a second event, *gsha1* underwent a tandem duplication to give rise to the near-by and highly similar *gsha2*. Under this scenario, we hypothesize that these three copies were kept throughout evolution as a mechanism to increase gene expression and protein production by increasing the number of gene copies within a single cell. This combined with somatic polyploidy would be a way to drastically increase the gene expression of a single cell.

The knockdowns of the two Hox genes *Scr* and *Ubx* resulted in the modification of the arolia into ectopic fans on T1 and T3 legs (Figure 3). It also demonstrated that the single giant cell which expresses *mogsha*/*gsha1*/*gsha2* and *CP19* is associated with the development of ectopic fans in those legs, reinforcing our claim that the giant cell drives the fan developmental program. In addition, this experiment suggests that the ancestral ground plan of the *Rhagovelia* lineage includes the fan developmental program in all three pairs of legs, and that the rewiring of *Scr* and *Ubx* has been necessary to block it in T1 and T3 legs. *Scr* and *Ubx* may have therefore connected, directly or indirectly, to the fan developmental program to repress fan formation in those legs. These results exemplify well how the emergence of innovative traits requires that developmental programs combine both new and pre-existing components by creating new interactions.

Finally, the silencing of *mogsha*/*gsha1*/*gsha2* expression using RNAi revealed that the giant cell was still present, suggesting that these genes do not play any role in the specification and differentiation of the giant cell. In addition, the RNAi did not cause *Ubx* to be expressed in the cells, indicating that other mechanisms restrict *Ubx* expression in the cell. Instead, our results suggest that *CP19* may be under the control of *mogsha*/*gsha1*/*gsha2* in young embryos. The effect of this downstream regulation of *mogsha*/*gsha1*/*gsha2* on *CP19* disappears in older embryos (Figure 4). It could be that *CP19* is co-regulated by other genes which take over to resume *CP19* expression in late embryonic stages. However, these results should be regarded with caution because the embryos were not precisely staged, and this hypothetical control of *CP19* may simply reflect differences in developmental stages between control and RNAi embryos. This experiment suggests that *mogsha*/*gsha1*/*gsha2* are mostly involved in the making of the fan, possibly by regulating cuticular genes, and the mechanisms responsible for the specification of the cell remain to be discovered.

In conclusion, our study sheds new light on the origin of key innovations. We indeed show that a single giant cell is sufficient for the development of the propelling fan in the *Rhagovelia* lineage. We also show that the emergence of this innovation requires that pre-existing Hox genes specify where in the embryo the giant cell and its new genetic program should develop. In addition, three paralogs, including two *Rhagovelia*-specific copies resulting from two successive duplication events, are necessary for the making of the fan. Taken together, the increase of gene copies via duplication and possibly via cell polyploidy may constitute a developmental solution to drastically increase the gene expression of a unique cell in order to produce and transport the material to make the fan. This study improves our understanding of the origin of key innovations, as it illuminates the cellular and developmental genetic basis of the fan, and reveals how a single cell can be sufficient to generate innovative traits that have been central for a lineage to access a new ecological niche and burst into diversification.

## Methods

### Animal rearing

We maintained a captive population of *Rhagovelia antilleana* in water tanks in a condition-controlled room (27°C; 55% humidity; 14h:10h light-dark cycles). Each water tank contained a bubbler, a water pump, plastic plants, and floating pieces of Styrofoam for females to lay eggs. All animals were fed daily with live and frozen house crickets.

### Library preparation, genome sequencing, and scaffolding

We generated a reference genome for *R. antilleana* using a combination of HiFi PacBio and Omni-C library preparation and sequencing, a hifiasm assembler, and a scaffolding of the assembly with Omni-C HiRise.

We flash-froze ca. 50 adult males and females in liquid nitrogen and shipped them to the Dovetail Genomics sequencing facilities. High Molecular Weight of genomic DNA was extracted and purified using a Qiagen Mini Kit (ref 13323) according the manufacturer’s instructions.

For PacBio library preparation and sequencing, DNA samples were quantified using Qubit 2.0 Fluorometer (Life Technologies, Carlsbad, CA, USA). The PacBio SMRTbell library (∼20kb) for PacBio Sequel was constructed using SMRTbell Express Template Prep Kit 2.0 (PacBio, Menlo Park, CA, USA) using the manufacturer recommended protocol. The library was bound to polymerase using the Sequel II Binding Kit 2.0 (PacBio) and loaded onto PacBio Sequel II). Sequencing was performed on PacBio Sequel II 8M SMRT cells.

PacBio CCS reads were used as an input to Hifiasm1 v0.15.4-r347 with default parameters. Blast results of the Hifiasm output assembly (hifiasm.p_ctg.fa) against the nt database were used as input for blobtools2 v1.1.1 and scaffolds identified as possible contamination were removed from the assembly (filtered.asm.cns.fa). Finally, purge_dups3 v1.2.5 was used to remove haplotigs and contig overlaps (purged.fa).

For each Dovetail Omni-C library, chromatin was fixed in place with formaldehyde in the nucleus. Fixed chromatin was digested with DNase I and then extracted, chromatin ends were repaired and ligated to a biotinylated bridge adapter followed by proximity ligation of adapter containing ends. After proximity ligation, crosslinks were reversed and the DNA purified. Purified DNA was treated to remove biotin that was not internal to ligated fragments. Sequencing libraries were generated using NEBNext Ultra enzymes and Illumina-compatible adapters. Biotin-containing fragments were isolated using streptavidin beads before PCR enrichment of each library. The library was sequenced on an Illumina Novaseq platform to produce ∼30x sequence coverage.

The input *de novo* assembly and Dovetail OmniC library reads were used as input data for HiRise, a software pipeline designed specifically for using proximity ligation data to scaffold genome assemblies (Putnam et al. 2016). Dovetail OmniC library sequences were aligned to the draft input assembly using bwa (https://github.com/lh3/bwa). The separations of Dovetail OmniC read pairs mapped within draft scaffolds were analyzed by HiRise to produce a likelihood model for genomic distance between read pairs, and the model was used to identify and break putative misjoins, to score prospective joins, and make joins above a threshold.

### Hybridization Chain Reaction – RNA fluorescence in situ hybridization (HCR-FISH)

We performed HCR-FISH to image mRNA expression patterns for multiple genes at the same time in embryos (Choi et al. 2018). We adjusted Bruce et al.’s protocol (Bruce et al. 2021) to our model species. First, to dissect embryos, we treated them with 50% bleach for 2:30 min, and removed the chorion and vitellin membranes with forceps in 1X PBS buffer and 0.05% Tween 20 (PTw 0.05%). Dissected embryos were then fixed in 4% paraformaldehyde (PFA) and heptane for 30 min, dehydrated in cold methanol 100% (washed 3 × 10 min), and placed at -20°C.

To start the HCR-FISH protocol, we rehydrated embryos in increasing concentrations of PTw 0.05% (3 × 10 min in MetOH 70%, 50%, and 30%, respectively) and performed successive washes in PTw 0.1% (4 × 10 min) to eliminate methanol. Embryos were then permeabilized using a detergent solution for 30 min (extended up to 2 h for older embryos with cuticle) and washed 10 min in PTw 0.1%. Embryos were then fixed again for 10 min in 4% PFA and washed in PTw 0.1% (3 × 10min). Embryos were pre-hybridized in pre-warmed probe hybridization buffer for 30 min à 37°C, then transferred in the probe solution (200 μL of probe hybridization buffer with 0.8 pmol of probe mixture obtained from Molecular Instruments) and left to incubate over night at 37°C.

The next day, we washed the embryos in pre-warmed probe wash buffer at 37°C (4 × 15 min) and in 5X SSCT at room temperature (2 × 5 min). Then, we pre-amplified embryos in amplification buffer for 30 min and, in the meantime, we activated the hairpin solution (2 μL of h1 and h2 in 100 μL of amplification buffer) with a heat shock (90 sec at 95°C and 30 min at room temperature), and we incubated the embryos in the hairpin solution overnight in the dark.

The next day, we removed the excess hairpins with five successive washes in 5X SSCT (2 × 10 min, 2 × 30 min, 1 × 10 min), and we incubated them in increasing concentration of glycerol (10 min in 30% glycerol, 10 min in 50% glycerol and 1μg/mL DAPI, and 10 min in 80% glycerol). Finally, we mounted the embryos between the slide and slip cover in 80% glycerol and observed using a confocal microscope (Zeiss LMS 780).

To quantify differences in gene expression based on our HCR-FISH imaging, we used ZEN Black 2.3. We compared the expression of *mogsha*/*gsha1*/*gsh2* and *CP19* in wild type and gsha-RNAi embryos. To do so, we used the polygon tool to select the giant cell, excluding the nucleus, and extracted the mean fluorescence intensity for the two genes in both conditions. This quantification method allows to compare relative gene expression for the same gene as long as the confocal imaging acquisition parameters were the same between samples, which was the case in our study.

### Nymphal RNA interference

We produced double stranded RNA (dsRNA) for *ubx* and *scr*. We designed forward and reverse primers both containing the T7 RNA polymerase promoter and used them to amplify the T7 PCR fragment using the previously cloned fragments as templates (see Santos et al. 2017 for primers information). The gene specific T7 PCR fragment was purified and used as a template to synthesize the dsRNA as mentioned in (Santos et al. 2017). We purified the resulting dsRNA using a RNeasy purification kit (Qiagen) and eluted it in Spradling injection buffer (Rubin and Spradling 1982) at a concentration of 5μg/μl. We injected fifth instar nymphs of *R. antilleana* with *scr* and *ubx* dsRNA, as described in (Khila et al. 2014). Injected nymphs were placed in water tanks, fed daily and allowed to develop until adulthood. We dissected and mounted adult legs and fans on slides in Hoyer’s medium, and observed them in an AxioObserverZ1 Zeiss widefield microscope.

### Maternal RNA interference

We followed the same procedure as described above to produce double stranded RNA (dsRNA) for *ubx, scr, gsha/mogsha*, except that we reached a dsRNA concentration of 7.5 μg/μl uing SpeedVac (Thermo Scientific). We injected adult females of *R. antilleana* with dsRNA of those genes and placed the females in water tanks along with adult males. We fed them daily and provided Styrofoam floaters for females to lay eggs on. We then collected the eggs at the relevant developmental stage and dissected them to perform an *in-situ* HCR. Some eggs were left to hatch to obtain phenocopies in first instar nymphs. We dissected and mounted the legs and fans of these nymphs on slides in Hoyer’s medium, and observed them in an AxioObserverZ1 Zeiss widefield microscope.

## Acknowledgements

We thank Ingrid Dourlens, Carlyne Golding, Juliette Mendes, Claudia Prûvot, Pascale Roux, and Mirjam Urb for their valuable advice and logistic help during the time of this study. This work was founded by Agence Nationale de la Recherche (ANR – project GEISHA).

