## Supplementary material for "A single giant cell at the origin of an evolutionary innovation": Supp Figure S1

**Supplementary Information Figure S1**

Images of adult phenotypes resulting from nymphal (4th instar) RNAi injections targeting *Scr* and *Ubx*. In *Scr*-RNAi individuals, the arolium normally present in wild type T1 legs (A) is replaced by an ectopic fan (B). In *Ubx*-RNAi individuals, fan integrity on T2 legs is not affected but the ventral claw loses its identity (E, H). On T3 legs, in contrast, the arolium normally present wild type individuals (C) is replaced by an ectopic fan (F).

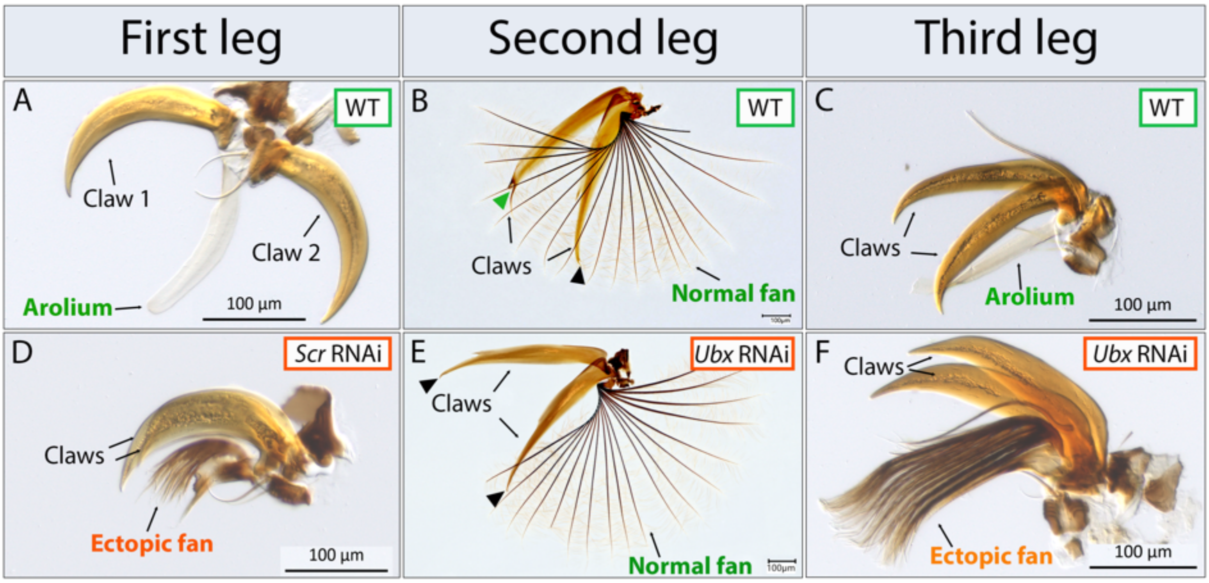
